# A drug repurposing screen reveals dopamine signaling as a critical pathway underlying potential therapeutics for the rare disease DPAGT1-CDG

**DOI:** 10.1101/2024.06.12.597770

**Authors:** Hans M. Dalton, Naomi J. Young, Alexys R. Berman, Heather D. Evans, Sydney J. Peterson, Kaylee A. Patterson, Clement Y. Chow

## Abstract

DPAGT1-CDG is a Congenital Disorder of Glycosylation (CDG) that lacks effective therapies. It is caused by mutations in the gene *DPAGT1* which encodes the first enzyme in N-linked glycosylation. We performed a drug repurposing screen on a model of DPAGT1-CDG in *Drosophila* (“*DPAGT1* model”) using 1,520 small molecules that are 98% FDA/EMA-approved. We identified 42 candidate drugs that improved the DPAGT1-CDG model. Notably from this screen, we found that pharmacological and genetic inhibition of the dopamine D2 receptor rescued the *DPAGT1* model. Loss of both dopamine synthesis and recycling rescued the model, suggesting that dopaminergic flux and subsequent binding to D2 receptors is detrimental under *DPAGT1* deficiency. This links dopamine signaling to N-glycosylation and represents a new potential therapeutic target for treating DPAGT1-CDG. We also genetically validate other top drug categories including acetylcholine-related drugs, COX inhibitors, and an inhibitor of NKCC1. These drugs and subsequent analyses reveal novel biology in DPAGT1 mechanisms, and they may represent new therapeutic options for DPAGT1-CDG.

## Introduction

Nearly 10% of the US population has a rare disease, yet over 95% lack effective therapeutics^1,2^. One solution to find new therapeutics is drug repurposing which use drugs that are already approved or under investigation^3^. When successful, drug repurposing screens provide immediate potential therapeutics that have already passed the rigor of human safety trials and may have a faster route of clinical approval^3^. Alternatively, they can potentially be used in an off-label capacity at a clinician’s discretion^3–5^. Even if a repurposed screen does not immediately result in a new therapy, it can still identify new biological interactions that can increase our understanding of a disorder.

DPAGT1-CDG is a rare disease that results from autosomal recessive loss-of-function mutations in the gene encoding the enzyme DPAGT1. It is one of nearly 200 Congenital Disorders of Glycosylation (CDGs)^6–9^. Glycosylation includes multiple pathways where sugars are co- or post-translationally added to proteins for their proper localization, folding, and function^10^. DPAGT1 is part of the N-glycosylation pathway where these sugars are added to specific asparagine (N) residues. DPAGT1 synthesizes dolichol-PP-GlcNAc which is the first step in N-glycosylation^6^.

DPAGT1-CDG causes developmental delay, muscle weakness, and seizures, among other symptoms^6,11,12^. Less severe mutations in *DPAGT1* can cause a form of congenital myasthenia syndrome (CMS) called DPAGT1-CMS^13^. Unlike DPAGT1-CDG, the CMS form has a known disease mechanism caused by hypoglycosylation of acetylcholine and calcium receptors^13,14^. Acetylcholinesterase inhibitors that increase acetylcholine levels can alleviate muscle weakness symptoms in both DPAGT1-CDG and -CMS patients^15–17^. However, there are still few treatment options for the multisystemic symptoms in DPAGT1-CDG.

Drug repurposing screens have had previous success in identifying new potential therapies for CDGs. For example, our laboratory identified GSK3β inhibitors as a potential treatment for NGLY1 deficiency^18^. In addition, a drug repurposing screen in *C. elegans* identified the aldose reductase inhibitor epalrestat for treating PMM2-CDG^19,20^. Thus, a drug repurposing screen for DPAGT1-CDG could help find new potential therapeutics for this disorder.

In this study, we performed a drug repurposing screen on a *Drosophila* model of DPAGT1-CDG to identify new therapeutics and biological interactions with DPAGT1. We identified 42 compounds that suppress the DPAGT1-CDG model and genetically confirmed many of the drug interactions. These 42 compounds include acetylcholine-related drugs, dopaminergic antagonists, and cyclooxygenase inhibitors, among others. Importantly, we find that both pharmacologic and genetic inhibition of the D2 receptor can rescue our *DPAGT1* model. In addition, impairing dopamine synthesis or recycling can also rescue the model. Our data suggest that manipulating dopamine signaling is a promising therapy for DPAGT1-CDG and implicates a larger role of dopamine signaling on N-glycosylation. The compounds and pathways identified in this screen are potential new therapeutic and genetic interactions with DPAGT1 that may represent new treatment options for DPAGT1-CDG.

## Results

### Identifying suppressors of a DPAGT1-CDG model in a primary drug repurposing screen

To model DPAGT1-CDG, we used an eye-based *Drosophila* model that we previously established^21^. In this model, *DPAGT1* expression is reduced in the eye by RNA interference (RNAi, UAS-GAL4 system^22^, *eya* composite-GAL4 driver^23^). This results in a small, improperly developed, rough eye phenotype (hereafter referred to as “*DPAGT1* model”)^21^ (Fig 1). The *DPAGT1* model can be chemically or genetically manipulated to make the eye larger (suppress the phenotype) or smaller (enhance the phenotype)^21^. To find potential therapeutics, we used the Prestwick Chemical Library, a collection of 1,520 compounds that are 98% FDA- or EMA-approved. We raised *DPAGT1* model flies on food containing each drug and measured the eye size of resulting progeny (Fig 1A). We used a 5 μM drug concentration - a standard dosage for flies that does not cause toxicity^18^. We scored 8,076 flies in total, and we calculated the Z-score of each compound by comparing their eye sizes to DMSO-treated *DPAGT1* model control flies. We identified 42 compounds that partially suppressed the eye phenotype of the *DPAGT1* model (“suppressors”) (Fig 1B and 1C, S1 and S2 Table) and 16 compounds that enhanced the eye phenotype (“enhancers”) (Fig 1B and 1C, S1 and S2 Table). 16 compounds resulted in no flies (S1 Table). Our positive hit rate of 2.8% falls in line with other compound screens performed in *Drosophila*^18,24–26^.

**Figure 1.**
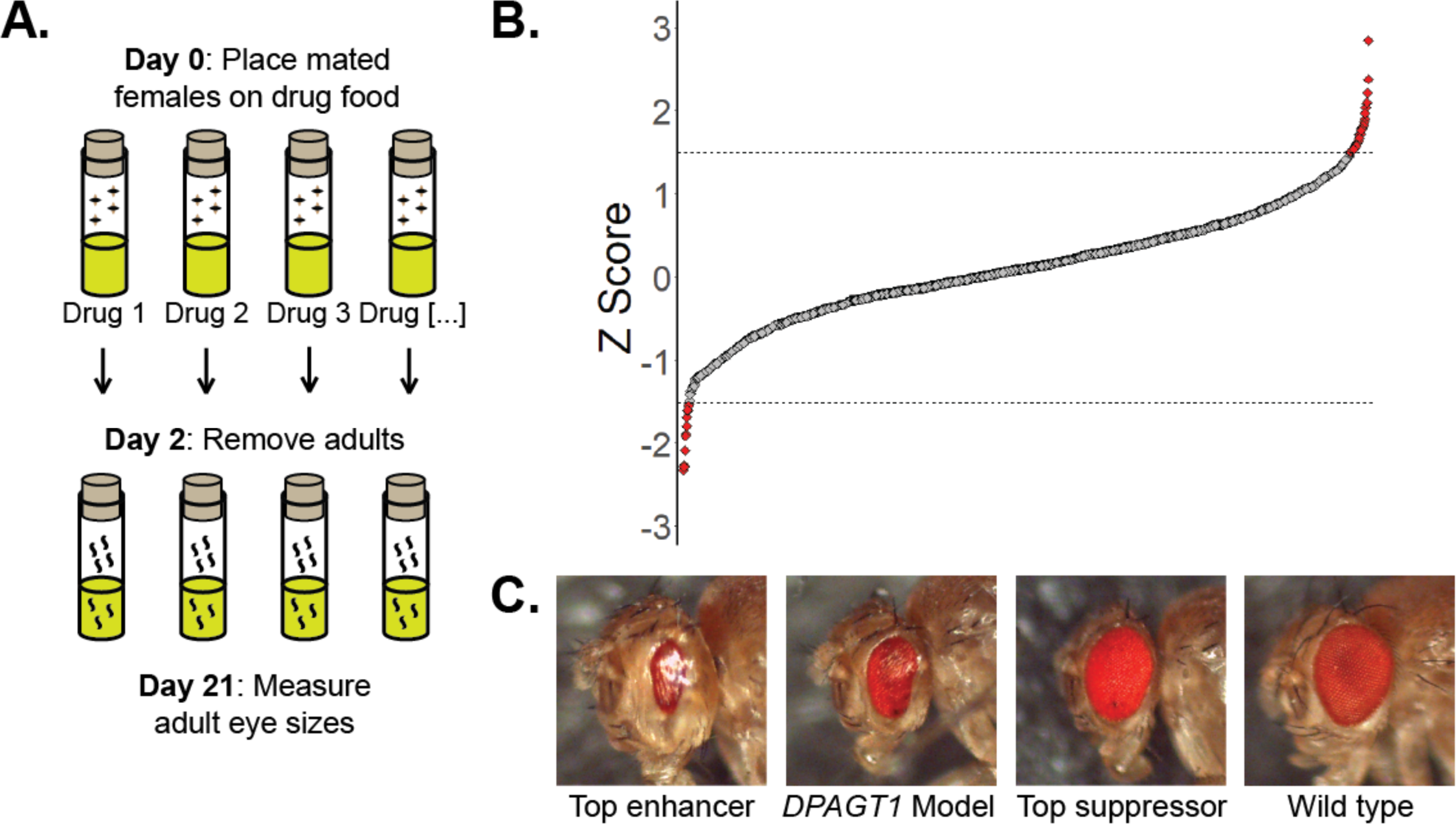
Summary of the drug repurposing screen. **(A)** Screening method for the drug repurposing screen. **(B)** Z-score plot of all repurposed drugs tested. Each point is the Z-score of a drug compared to the average of DMSO-treated control flies. Dotted lines indicate our Z-score threshold of 1.5 or -1.5. We found 42 and 16 drugs with a Z-score of ≥1.5 or ≤-1.5, respectively. **(C)** Representative images showing the effects of the top enhancer and suppressor, the *DPAGT1* model, and a control eye containing the *eya* composite-GAL4 driver alone. See Table S1 for a complete list of compounds and Z-scores.

We focused our validation efforts on suppressors that had ≥3 drugs in a particular drug class, had the strongest suppressive effect on eye size (Z-score ≥2), or had high potential for patient use (e.g. available over-the-counter). Drug classes with the highest representation were acetylcholine-related drugs, dopamine receptor antagonists, COX inhibitors, and antioxidants (three drugs each, S2 Table). In total, we tested 20/42 (48%) of the suppressors in validation experiments.

### Impairing acetylcholine breakdown improves the *DPAGT1* model

Mutations in *DPAGT1* can cause muscle weakness because acetylcholine receptors (AChRs) become hypoglycosylated^13,14^. This muscle weakness can be treated by cholinesterase inhibitors which increase available acetylcholine^13,14,27^. Our unbiased screen identified three AChR-related drugs. This included methacholine, an analog of acetylcholine and a muscarinic receptor agonist^28^, as well as two drugs expected to increase acetylcholine levels: benactyzine, a cholinesterase inhibitor/anticholinergic^29^, and neostigmine, a cholinesterase inhibitor^30^. To validate drug hits from the screen, we primarily used drugs purchased from a second source to reduce potential quality control errors (this is true of all further validation) (S2 Table). In dose-response analyses, benactyzine and neostigmine both increased the eye size in the *DPAGT1* model (Fig 2A and 2B). Benactyzine significantly increased female eye size at 25 μM and generally increased the upper distribution of eye sizes at most doses tested in both sexes (Fig 2A). Neostigmine significantly increased eye size in males at 5 and 25 μM (Fig 2B) and had a trending (+12%), but not statistically significant, increase in median size in females at 25 μM. As acetylcholinesterase inhibitors are already used in patients, these two drugs represent a strong validation of the screen to find patient-relevant hits.

**Figure 2.**
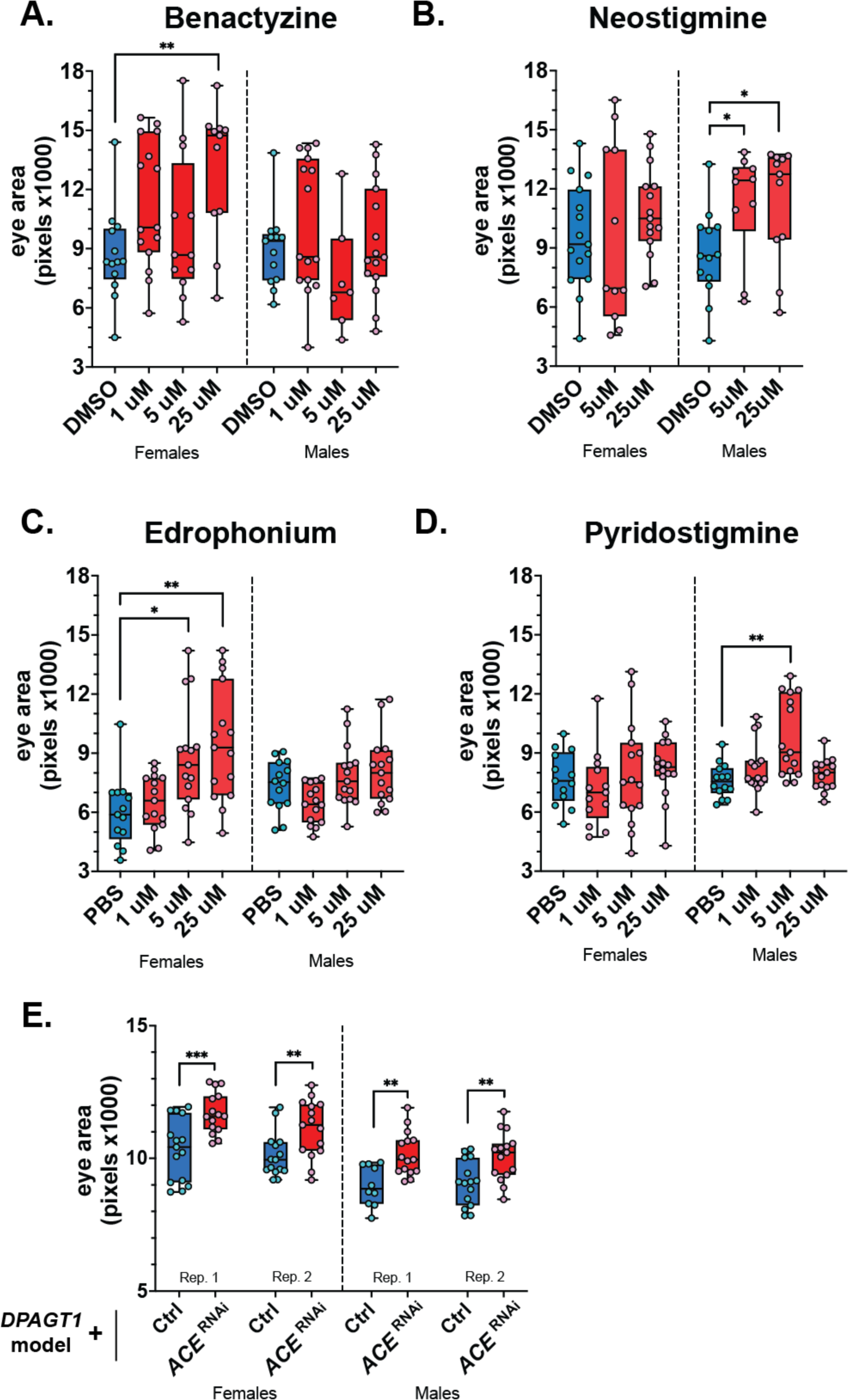
Pharmacological and genetic inhibition of acetylcholinesterase improved the *DPAGT1* model. **(A-D)** Multiple ACh-related drugs can partially rescue the *DPAGT1* model. For Neostigmine, we assayed 1 μM separately, but it was not significant. (**E)** RNAi knockdown of *ACE* also partially rescues the *DPAGT1* model (VDRC 105432KK). See S2 and S3 Table for more details. * p<0.05, ** p<0.01, *** p<0.001 (Student’s t-test or One-way ANOVA with multiple comparison correction).

We next assayed four acetylcholine-related drugs already used by DPAGT1-CDG or -CMS patients for their effect on the *DPAGT1* model. This included pyridostigmine (used in DPAGT1-CMS^15,17^ and DPAGT1-CDG patients^16^), amifampridine, salbutamol, and edrophonium (all used in DPAGT1-CMS patients, though edrophonium is more diagnostic^15,31^). Treatment with edrophonium strongly improved female *DPAGT1* model flies in a dose-dependent manner (Fig. 2C). In males, while not significantly changing the median eye size, edrophonium treatment did cause several positive outliers at 5 and 25 μM (Fig. 2C). Similar to its analog neostigmine (Fig 2B), treatment with pyridostigmine caused a significant increase in eye size in males at 5 μM (Fig. 2D) and a small, but not significant increase in females at 25 μM (+6%). Overall, acetylcholine-affecting drugs identified in the screen and currently used by patients improved the *DPAGT1* model.

The rescuing drugs neostigmine, pyridostigmine, and edrophonium all inhibit the enzyme acetylcholinesterase (AChE) which breaks down acetylcholine^14^. We tested whether genetic inhibition of the AChE-encoding gene *ACE* (human: *ACHE*) could rescue the model as well. We used two RNAi lines against *ACE* (via the GAL4/UAS system^22^) to mimic pharmacological inhibition. One previously studied^32^ RNAi line improved the *DPAGT1* model in both sexes (Fig. 2E), mimicking the drug inhibition of *ACE*. The second RNAi line was capable of improving females (S3 Table). There was little effect from *ACE* knockdown on its own (S3 Table). These pharmacological and genetic data indicate that inhibition of AChE, and the ostensible increase in acetylcholine, can partially rescue the eye development defect of the *DPAGT1* model. This validates the screen, supports continued usage of AChE inhibitors in patients, and might also indicate a role for AChE inhibitors outside of their effect on muscle weakness.

### Antagonizing D2 receptor signaling significantly improves the *DPAGT1* model

Dopamine (DA) synthesis, transport, and downstream receptor signaling is well-conserved between *Drosophila* and humans^33,34^ (Fig. 3A). We will primarily discuss female data here, as females had an overall more robust response to manipulating dopamine signaling. All results, including male data, can be found in S2 and S3 Tables. Three DA receptor antagonists were suppressors with moderate Z-scores in our screen (Z-scores: 1.8-1.86, S1 Table). These included sulpiride, a D2/D3 receptor antagonist^35^, paliperidone, an antipsychotic that antagonizes D2 and 5-HT2A receptors^36^, and prochlorperazine, a first-generation antipsychotic that primarily antagonizes D2 receptors but also targets adrenergic, cholinergic, and histaminergic receptors^37–39^. Of these, secondary testing of prochlorperazine improved the *DPAGT1* model at 1 μM in females (Fig 3B), and paliperidone also improved male flies (S2 Table). While evaluating our drug classes, we observed that one D2 receptor antagonist - trifluoperazine^40^ - was excluded for technical reasons. We retested this drug and found a robust increase in eye size in the *DPAGT1* model at the 5 μM dose in both sexes (Fig 3C, S2 Table). While prochlorperazine, paliperidone, and trifluoperazine overlap in function on D2 receptors, they can also target other receptors and pathways. Given this, we focused on DA signaling by genetically reducing the expression of their shared target *Dop2R* (human: *D2R*) in the *DPAGT1* model.

**Figure 3.**
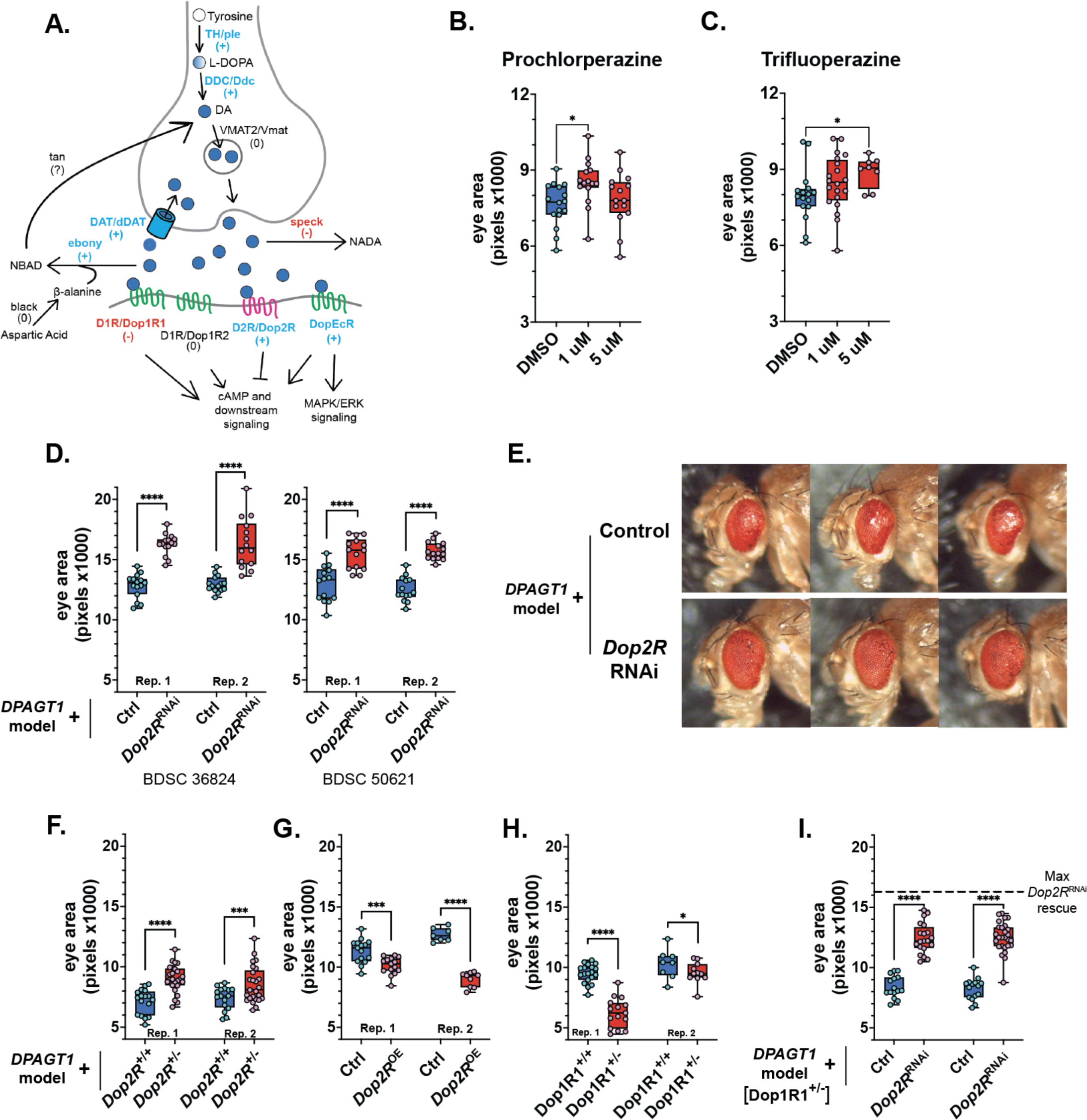
Inhibiting the dopamine 2 receptor improved the *DPAGT1* model. (**A)** A schematic of dopamine synthesis, recycling, metabolism, and signaling in *Drosophila*. The effect of each RNAi or mutation on the *DPAGT1* model is indicated. Genes in blue with the “+” symbol improved the *DPAGT1* model when knocked down, while those in red with the “-” symbol made it worse. Genes in black with the “0” symbol had no effect or could not be distinguished from the knockdown effect on its own. **(B-C)** The D2 antagonists prochlorperazine and trifluoperazine improved eye size in the *DPAGT1* model. For technical reasons, we assayed 25 μM separately, but neither drug was significant. **(D)** RNAi knockdown of the *Dop2R* gene (BDSC 36824) improved the *DPAGT1* model. **(E)** Representative images comparing the *DPAGT1* model crossed with the control RNAi background (attP2, BDSC 36303) or a *Dop2R* RNAi line (BDSC 36824). **(F)** *DPAGT1* model flies carrying a heterozygous null of *Dop2R* had increased eye size (BDSC 84720). **(G)** *DPAGT1* model flies crossed to a *Dop2R* overexpression line (UAS-Dop2R, BDSC 86134) had worse eye size. **(H)** A heterozygous null of *Dop1R1* (BDSC 92640) results in worse eyes in the *DPAGT1* model. **(I)** *Dop2R* RNAi (BDSC 36824) improved *DPAGT1* model [*Dop1R1^+/-^*] flies, but to a smaller maximal eye size than in *DPAGT1* model flies alone (indicated by dashed line, averaged from Fig 3D). All graphs are of female flies. See S2 and S3 Table for more details. * p<0.05, *** p<0.001, **** p<0.0001 (Student’s t-test or One-way ANOVA with multiple comparison correction).

*Dop2R* is the only *Drosophila* ortholog of human genes *D2R*, *D3R*, and *D4R*. These encode for “D2-like” G-protein coupled receptors (GPCRs) that inhibit the adenylate cyclase/cAMP signaling pathway^33,34^ (Fig 3A). Matching the pharmacological data from the primary screen, reduced expression of *Dop2R* using RNAi resulted in a strong improvement of eye size across all RNAi lines and replicates tested (Fig 3D and 3E, S3 Table). In addition, their eyes showed strong qualitative improvement in ommatidium structure with fewer “glassy” eye sections (Fig 3E). We also crossed a heterozygous *Dop2R* null allele^41^ into the *DPAGT1* model (note that *Dop2R* is X-linked and only females are analyzed here). Mimicking the RNAi, *DPAGT1* model flies carrying a heterozygous null *Dop2R* mutation also resulted in improved eye size (Fig 3F). This indicates that this improvement is not due to any indirect UAS-GAL4^22^ effects. Finally, overexpressing *Dop2R*^41^ in the *DPAGT1* model decreased eye size (Fig 3G), confirming what is expected given the knockdown and null result. Genetically manipulating *Dop2R* had little effect on its own (S3 Table). Overall, this indicates that reduction of D2 receptor signaling improves the *DPAGT1* model.

After release into the synaptic cleft, DA can bind to several receptors in the fly. This includes the Dop2R receptor as well as the Dop1R1, Dop1R2, and DopEcR receptors^33,34^. *Dop1R1* and *Dop1R2* are orthologs of the human *D1R* and *D5R* genes. These encode “D1-like” GPCRs that activate the adenylate cyclase/cAMP signaling pathway^33,34^ (opposite of “D2-like” receptors). Given the opposite downstream effect of these receptors compared to *Dop2R*, we hypothesized that they would have an opposite effect on eye size. We crossed heterozygous null mutations of *Dop1R1*^42^ and *Dop1R2*^41^ into the *DPAGT1* model. Supporting our hypothesis, *DPAGT1* model flies carrying a heterozygous null *Dop1R1* mutation had worse eyes than the *DPAGT1* model alone (Fig 3H, S3 Table). *DPAGT1* model flies carrying a heterozygous null *Dop1R2* mutation had only a mild effect in one female replicate and no effect in males (S3 Table).

Given the strong rescue effect of *Dop2R* RNAi (Fig 3D-E), we tested whether this loss of *Dop2R* could rescue the loss of *Dop1R1* in the *DPAGT1* model. We created a recombinant line of the *DPAGT1* model with the *Dop1R1* null mutation (*DPAGT1* model [*Dop1R1^+/-^*]). Loss of *Dop2R* does rescue the *DPAGT1* model [*Dop1R1^+/-^*] eye size, but to a lesser extent than loss of *Dop2R* on its own (Fig 3I, p<0.0001, Student’s t test, *Dop2R* RNAi compared across experiments). This result is in line with the opposing effects of *Dop1R1* and *Dop2R* on downstream cAMP signaling (Fig 3A).

The fourth DA receptor in *Drosophila*, DopEcR, has no clear human ortholog. Similar to Dop1R1, DopEcR activates the downstream adenylate cyclase/cAMP signaling pathway through DA binding. However, DopEcR can also activate the Mitogen-Activated Protein Kinase (MAPK) pathway through binding of the insect steroid ecdysone^33,34^. RNAi knockdown of *DopEcR* also improved eye size in the *DPAGT1* model (S3 Table). Taken together, these data suggest that loss of DA signaling through Dop2R and DopEcR is beneficial to the *DPAGT1* model, while loss of DA signaling through Dop1R1 is detrimental to the model.

### Impairing dopamine synthesis and recycling can improve the *DPAGT1* model

DA is synthesized by the enzymes tyrosine hydroxylase and DOPA decarboxylase, which are encoded by the *ple* and *Ddc* genes, respectively (humans: *TH* and *DDC*). DA is then transported into vesicles by the vesicular monoamine transporter VMAT, encoded by *Vmat* (humans: *VMAT2*), before being shuttled into the synaptic cleft^33,34^ (Fig 3A). Similar to knockdown of *Dop2R*, knockdown of the first DA synthesis gene, *ple*, and the second DA synthesis gene, *Ddc,* strongly improved the *DPAGT1* model (Fig 4A and 4B, S3 Table). Knockdown of *Vmat* also improved the model, but it had a similar increase on its own (S3 Table). Overall, this suggests that inhibiting DA synthesis is beneficial to the *DPAGT1* model.

**Figure 4.**
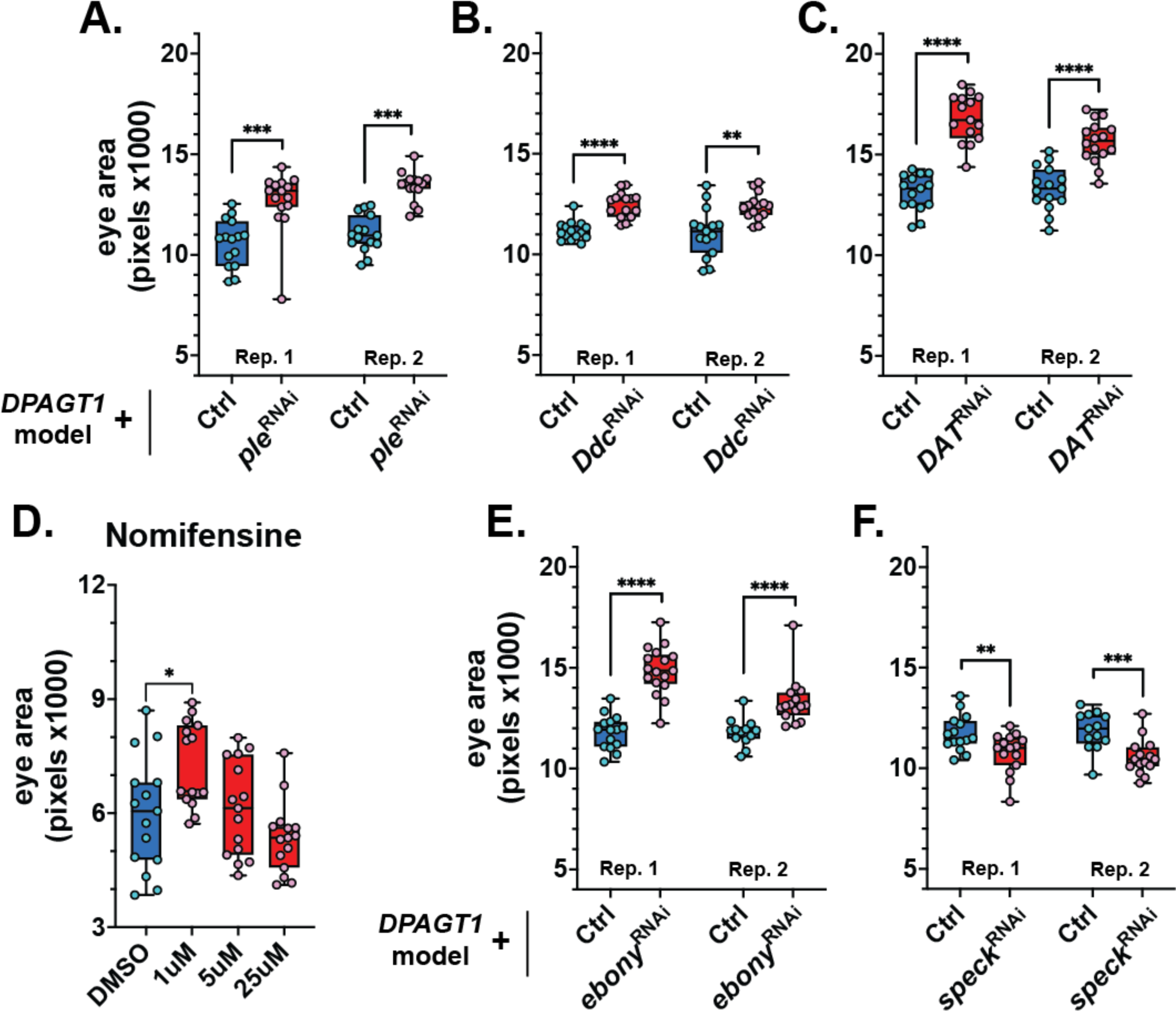
Inhibiting dopamine synthesis and recycling improves the *DPAGT1* model. **(A-B)** RNAi targeting the genes encoding the DA synthesizing enzymes ple and ddc (BDSC 25796 and 27030) improved the *DPAGT1* model. **(C-D)** RNAi against the gene encoding the DA recycling protein DAT (BDSC 50619), or pharmacologic inhibition of DAT via nomifensine at 1 μM, improved the *DPAGT1* model. **(E)** RNAi against the gene encoding the DA recycling protein ebony (BDSC 28612) improved the *DPAGT1* model. **(F)** RNAi against the gene encoding the N-acetyltransferase speck resulted in worse eyes in the *DPAGT1* model. All data are of female flies. See S2 and S3 Table for male data and more details. * p<0.05, ** p<0.01, *** p<0.001, **** p<0.0001 (Student’s t-test or One-way ANOVA with multiple comparison correction).

There are several ways that DA can exit the synaptic cleft. DA can be recycled back from the synaptic cleft to the presynaptic neuron by the DA transporter DAT^33,34^ (encoded by *DAT*) (Fig 3A). Knockdown of *DAT* strongly improved *DPAGT1* model eye size, matching the same increase as loss of *Dop2R* (Fig. 3C, Fig 4C, S3 Table). The DAT inhibitor^43,44^ nomifensine maleate had a positive Z-score in the primary screen (0.84) but did not reach our hit threshold. Because we saw strong improvements in eye size when *DAT* was knocked down, we tested a version of this drug lacking the maleate salt (S2 Table). In line with the genetic data, treatment with nomifensine improved the *DPAGT1* model at 1 μM (Fig 4D). In *Drosophila,* but not in humans, DA can also be converted to the metabolic product NBAD for translocation and conversion back into DA. These steps are performed by the enzymes black, ebony, and tan through the β-alanylation pathway^33,34^ (encoded by *black*, *ebony*, and *tan*) (Fig 3A). ebony and tan convert and shuttle DA, while black creates the β-alanine necessary for ebony to function^33^. RNAi knockdown of the first β-alanylation gene, *ebony*, strongly improved the *DPAGT1* model (Fig 4E, S3 Table). However, while knockdown of *black* improved the *DPAGT1* model, it had a similar increase on its own (S3 Table). There are no available RNAi lines for *tan*. Overall, inhibiting the recycling of DA back into the synaptic cleft through DAT or ebony improves the *DPAGT1* model.

In contrast to these recycling pathways, DA can exit the synaptic cleft through metabolism into the inactive compound NADA in *Drosophila*. This occurs through the N-acetyltransferase enzyme, speck, and prevents further signaling by that DA molecule. While speck has no clear human ortholog, it serves a similar role to monoamine oxidases in humans^33^. Because speck removes DA, we hypothesized that speck would have the opposite effect of the DA synthesis genes. Supporting this hypothesis, knockdown of *speck* resulted in worse eye sizes in the *DPAGT1* model (Fig 4F, S3 Table). Thus, inhibiting the removal of DA from the synaptic cleft is detrimental to the *DPAGT1* model.

Generally, these results fit the hypothesis that most *DPAGT1* model outcomes can be predicted by how a treatment affects DA binding to Dop2R (Fig 3A). For example, inhibiting DA synthesis or recycling may improve the *DPAGT1* model by reducing the flux of DA to Dop2R. In contrast, because speck normally removes DA and reduces its binding to Dop2R, this could explain why inhibiting speck worsens the *DPAGT1* model. Overall, these data strongly link the inhibition of DA synthesis, recycling, and signaling with improvement of *DPAGT1* deficiency. Drugs that impair D2 signaling or synthesis, or are agonists of D1 receptors, may be good therapeutics for DPAGT1-CDG.

### Histaminergic signaling is beneficial to the *DPAGT1* model

The Histamine 2 (H2) receptor antagonist ranitidine, used to decrease stomach acid^45^, was a strong enhancer in our screen (Z-score = -2.33, S1 Table). Histamine and dopamine have some overlapping biology. For example, *Drosophila* can process dopamine and histamine using the same β-alanylation enzymes - ebony, black, and tan^33,46,47^. In addition, loss of histamine can upregulate depolarization-stimulated DA release and receptor expression in mice^48^. Given the impact of DA signaling on the *DPAGT1* model (Fig 3 and 4), our results from *ebony* knockdown (Fig 4E, S3 Table), and the finding of ranitidine, we tested histaminergic signaling further. In line with the primary screen, ranitidine significantly worsened eye size in the *DPAGT1* model at multiple doses (Fig 5A). As ranitidine is an H2 receptor antagonist, we tested if exogenous histamine (an H1/H2 agonist) would have the opposite effect. Treatment with histamine showed statistically significant increase at 5 μM in males and increased the upper distribution of both sexes at 25 μM (Fig 5B). Histamine is an atypical medication which is used in select cancer therapies^49^. Histamine could represent a new therapeutic for patients, and these data also suggest that some antihistamines may be detrimental under *DPAGT1* impairment.

**Figure 5.**
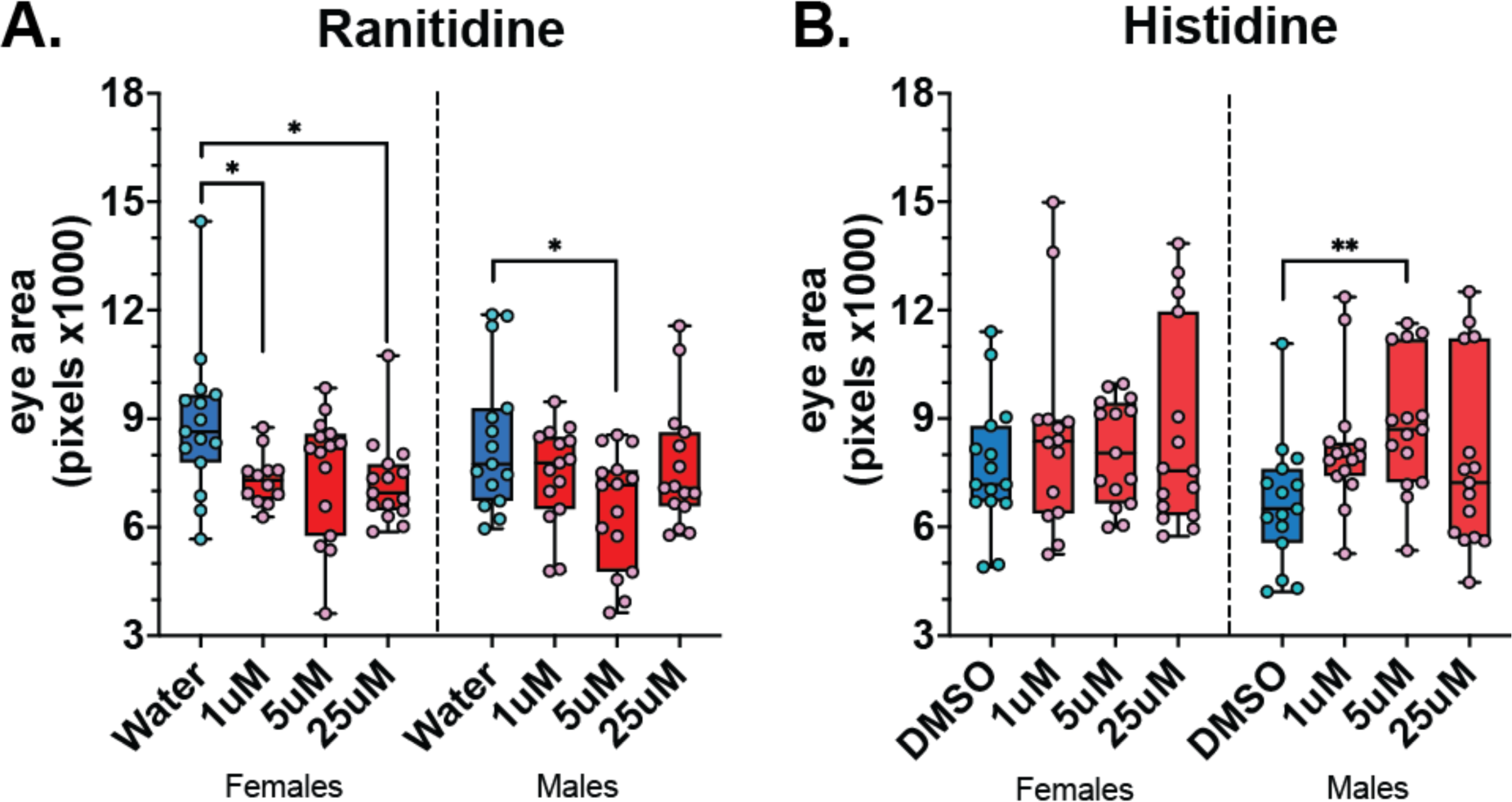
Antagonizing histamine signaling worsens the *DPAGT1* model. **(A)** The H2 antagonist Ranitidine worsens the *DPAGT1* model at multiple concentrations. **(B)** The H1/H2 agonist histamine can partially rescue the *DPAGT1* model at 5 μM in males. See S2 Table for more details. * p<0.05, ** p<0.01 (One-way ANOVA with multiple comparison correction).

### Inhibiting the ion transporter NKCC1 improves the *DPAGT1* model

The loop diuretic bumetanide was a strong suppressor hit in our screen (Z = 2.04). Bumetanide inhibits the ion cotransporters NKCC1 and NKCC2 (encoded by *NKCC1* and *NKCC2* in humans). Both proteins transport Na+, K+, and Cl-ions into cells primarily in the secretory epithelia or renal tissues, respectively^50^. In addition to its diuretic property, bumetanide has recently been tested as an off-label anti-seizure medication to some success^51–54^. Finally, we previously found that NKCC1 is a genetic modifier of NGLY1 deficiency, another CDG^55^.

Matching the primary screen, bumetanide caused a strong increase in eye size in the *DPAGT1* model in a dose-dependent manner (Fig 6A). *Ncc69* is the *Drosophila* ortholog of *NKCC1* and *NKCC2*^55,56^. Genetic knockdown of *Ncc69* also resulted in an increase in eye size in the *DPAGT1* model (Fig 6B, S3 Table). This corroborates with the bumetanide data that inhibiting NKCC1/NKCC2 activity is beneficial under *DPAGT1* deficiency. Thus, drugs that inhibit NKCC1 activity may be beneficial when *DPAGT1* activity is reduced.

**Figure 6.**
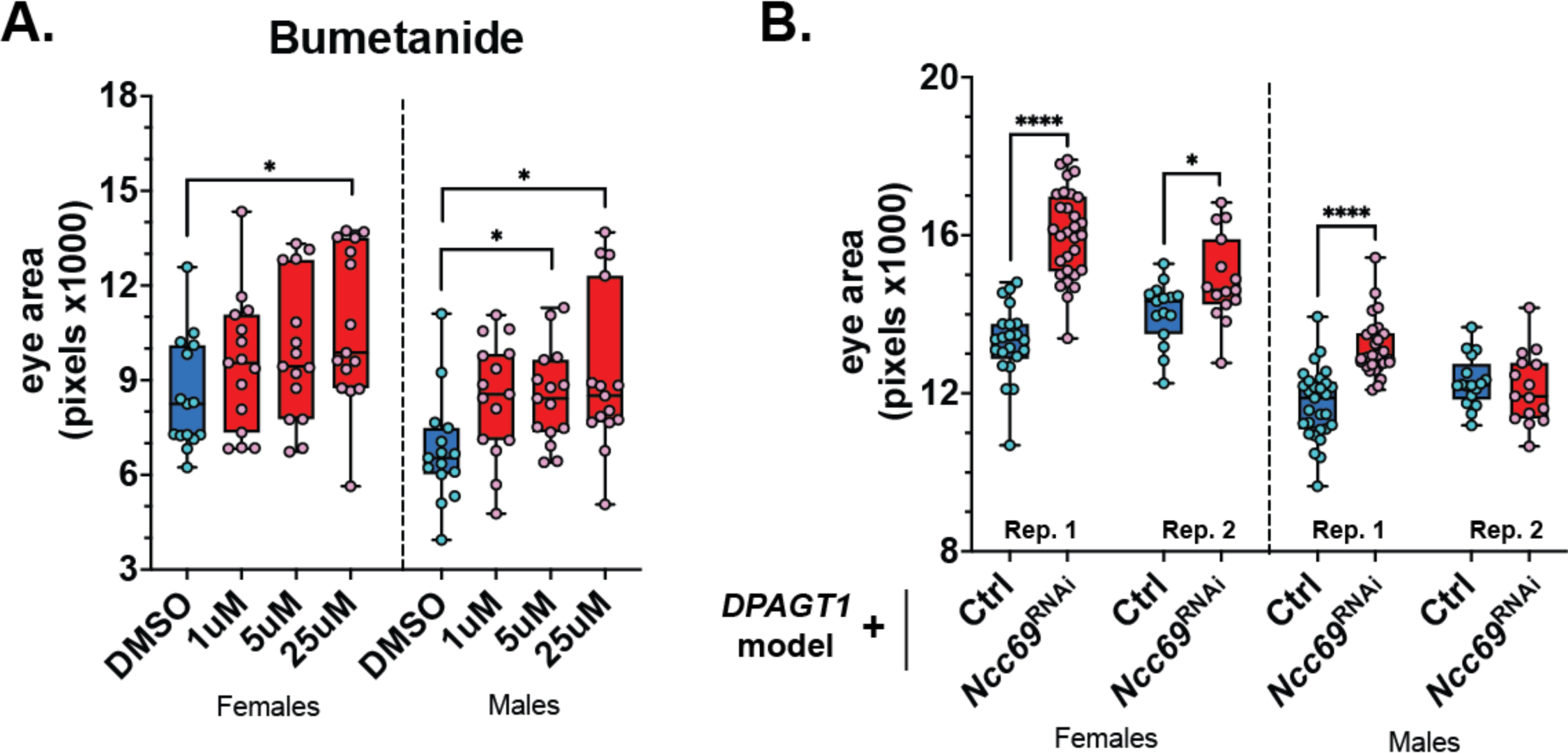
Inhibiting the ion transporter NKCC1 improves the *DPAGT1* model. **(A)** Bumetanide increases eye size in the *DPAGT1* model. **(B)** Genetic knockdown of *Ncc69*, which encodes the target of bumetanide inhibition, also increases eye size in the *DPAGT1* model. See S2 and S3 Table for more details. * p<0.05, **** p<0.0001 (Student’s t-test or One-way ANOVA with multiple comparison correction).

### COX inhibitors improve the *DPAGT1* model

The cyclooxygenase signaling pathway involves the synthesis of prostaglandins that are important for processes such as inflammation and growth^57–59^. Many non-steroidal, anti-inflammatory drugs (NSAIDs), such as ibuprofen, inhibit the first enzymes in this process - cyclooxygenases COX-1 and COX-2 (encoded by *COX1* and *COX2* in humans). There were three COX-1/COX-2 inhibitor hits in our screen: triflusal^60^, antipyrine, and the antipyrine metabolic product, 4-hydroxyantipyrine^61^. Of these, antipyrine significantly increased average eye size at the 1 μM dose, and multiple antipyrine doses resulted in a positive shift in *DPAGT1* model eye size distributions (Fig 7A).

**Figure 7.**
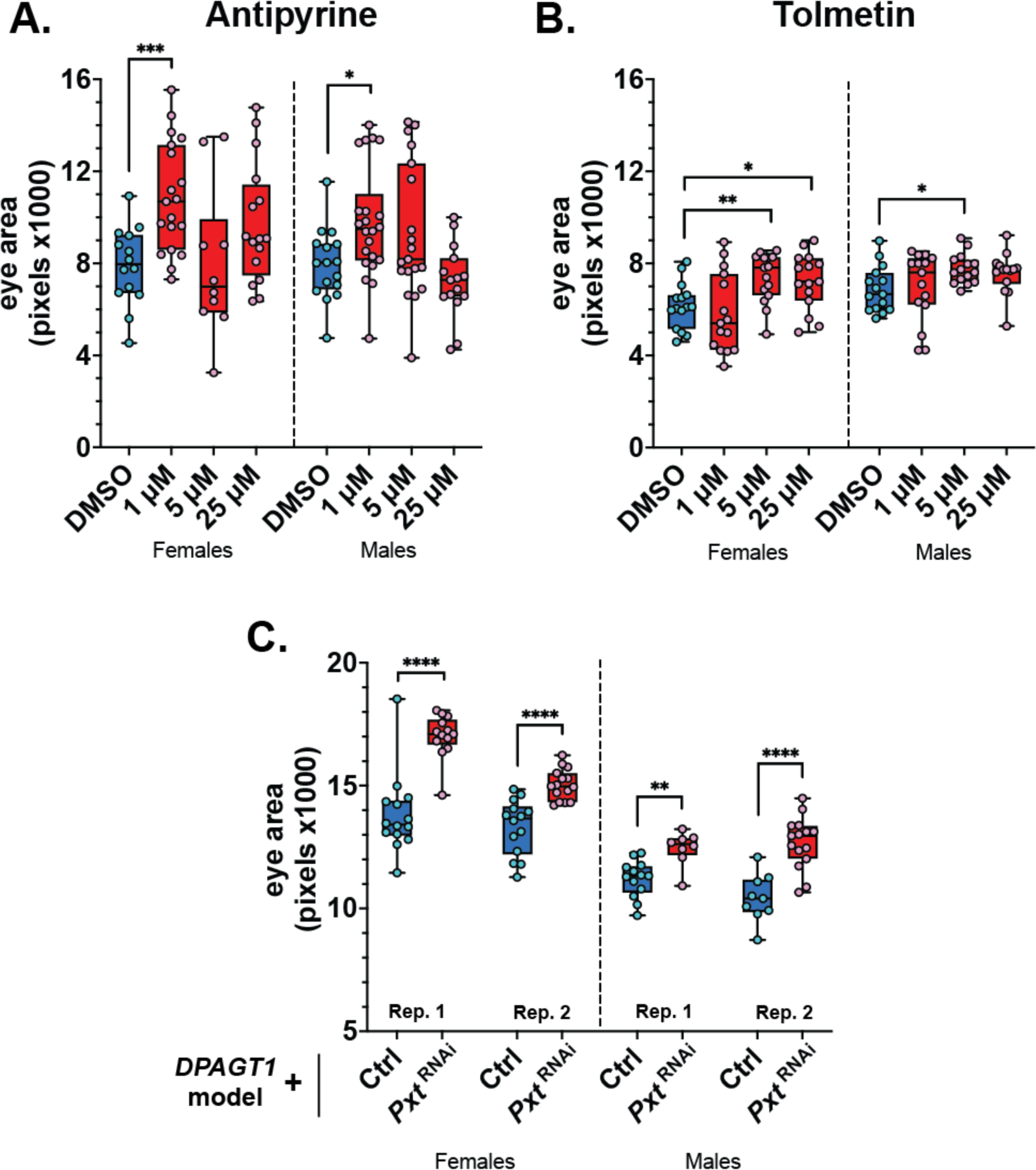
Inhibiting COX enzymes improves the *DPAGT1* model. **(A-B)** Both antipyrine and tolmetin improved the eye size of the *DPAGT1* model. **(C)** RNAi against the gene encoding the COX-like enzyme Pxt (BDSC 32382) partially rescued the *DPAGT1* model. See S2 and S3 Table for more details. * p<0.05, ** p<0.01, ** p<0.001, **** p<0.0001 (Student’s t-test or One-way ANOVA with multiple comparison correction).

NSAIDs are commonly used and would be easy to access for patients^62^. However, COX inhibitors can vary in their specificity and degree of COX inhibition, and this can affect their efficacy and tolerance^63–66^. As such, we tested five other COX inhibitors using the *DPAGT1* model. Of these, the COX-1/COX-2 inhibitor tolmetin^67^ significantly improved the *DPAGT1* model at 5 μM in both sexes, and at 25 μM in females (Fig 7B). The *Drosophila* gene *Pxt* is the ortholog of human *COX1* and *COX2*^68,69^. Similar to antipyrine and tolmetin, knockdown of *Pxt* also resulted in improved eye size in the *DPAGT1* model (Fig 7C, S3 Table). Given that inhibiting *Pxt* is beneficial, we hypothesize that antipyrine and tolmetin specifically improve the *DPAGT1* model because they may be more efficacious or well-tolerated under *DPAGT1* deficiency. Overall, inhibiting prostaglandin signaling through COX inhibition is beneficial to the *DPAGT1* model, and certain COX inhibitors may be useful in alleviating symptoms of *DPAGT1* impairment in patients.

## Discussion

Here, we identify multiple drugs that improve a *Drosophila* model of *DPAGT1* deficiency. Overall, 8/21 (38%) drugs tested from the primary screen validated in further dose-response analysis. In addition, 6/16 (38%) drugs derived from the primary screen improved the *DPAGT1* model. Almost all tested drugs were genetically validated by manipulating their target genes in *Drosophila*. We found that drug classes involving acetylcholine signaling, dopamine synthesis, histamine receptors, NKCC1/2, and cyclooxygenase inhibition might be good candidates for treating DPAGT1-CDG.

One of the enriched classes was acetylcholine (ACh)-related drugs. Identifying this drug class provides strong validation of our fly model and drug screen, as this class is already used by both DPAGT1-CDG and -CMS patients^15,16,31^. In addition, multiple AChE inhibitors currently taken by patients improved the *DPAGT1* model. This indicates that our CDG fly model is a good representation of the disorder, and that these models are valuable tools for identifying potential therapeutics. We found that genetically or pharmacologically inhibiting acetylcholine breakdown is beneficial under *DPAGT1* impairment. Acetylcholinesterase inhibitors are taken to improve muscle function in *DPAGT1* patients by increasing acetylcholine signaling at neuromuscular junctions (NMJs)^70^. However, in *Drosophila*, glutamate is the excitatory neurotransmitter in NMJs^71^, and the fly eye is likely not impacted by NMJ signaling regardless. Outside of NMJs, acetylcholine is important for many biological processes, including neurodevelopment^72–74^ and growth^75–77^. Given the impaired eye development in the *DPAGT1* model, these processes may underlie the improvement from *ACE* inhibition. Acetylcholine can also bind to nicotinic acetylcholine receptors to mediate dopamine (DA) release in flies^78^ and mammals^79,80^. Taken together with our results in the DA signaling pathway, this could indicate an overlapping mechanism. Overall, acetylcholinesterase inhibitors have beneficial effects under *DPAGT1* impairment. These drugs are worth exploring further in patients, beyond their role in improving muscle weakness, given the improvement in eye development in our model.

We found that inhibiting Dop2R signaling improves the *DPAGT1* model. In addition, knockdown of DA synthesis or recycling genes improves the model. Thus, the synthesis of DA, and its binding to the Dop2R, are detrimental under *DPAGT1* inhibition. Dop2R signaling inhibits the downstream adenylate cyclase/cAMP signaling pathway. While cAMP signaling has many diverse functions, it is important for proper neuronal connectivity and survival^81,82^. Therefore, inhibiting Dop2R signaling may increase cAMP signaling and allow for improved neuronal outcomes. This could underlie the benefits in the *DPAGT1* model from Dop2R genetic and pharmacological inhibition. Supporting this hypothesis, heterozygous knockout of the Dop1R1 receptor worsened the *DPAGT1* model. Binding to Dop1R1 may counteract signaling to Dop2R through increased cAMP signaling. Given this, drug classes such as D2R antagonists or D1R agonists might be good therapeutics for DPAGT1-CDG. Of note, we identified two monoamine oxidase inhibitors (MAOIs) as enhancers in the screen (moclobemide and selegiline, S2 Table). MAOIs inhibit MAOs to prevent the breakdown of neurotransmitters such as DA^83^. It is possible that these MAOIs were enhancers because they increased the amount of DA in the presynaptic neuron, in line with our data on DA synthesis and recycling.

Misregulated DA signaling is associated with neurological disorders such as Parkinson’s disease, schizophrenia, and bipolar disorder^84^. DPAGT1-CDG has no previous clear connection to DA. However, both DPAGT1-CDG and -CMS patients have abnormal gait^31,85^. DPAGT1-CDG patients also have neurological symptoms including seizures and intellectual disability^6,12^. Misregulated DA could underlie some symptoms of DPAGT1-CDG. Interestingly, the DA pathway is misregulated in a fly model of another CDG, NGLY1 deficiency^18^. Further, aripiprazole, a complex agonist of D2 receptors^86^, could rescue worm and fly disease models of NGLY1 deficiency^87^. Thus, DA signaling may be important in glycosylation disorders more generally.

Inhibiting the ion transporter Ncc69 via bumetanide or RNAi resulted in a strong improvement of the *DPAGT1* model. In humans, NKCC1 has 3 sites of N-glycosylation while NKCC2 has 2 (via Uniprot^88^), and inhibiting N-glycosylation reduces the function of these enzymes^89,90^. In addition, we previously found that loss of the deglycosylating enzyme NGLY1 results in reduced NKCC1 activity^55^. Given this, impaired N-glycosylation in the *DPAGT1* model likely reduces the function of Ncc69. Thus, it was surprising that further reducing its expression by RNAi actually improved the *DPAGT1* model. We hypothesize that under *DPAGT1* impairment, Ncc69 may be misglycosylated and subsequently may be misfolded, or have aberrant function, and contribute to disease pathogenesis.

Bumetanide is typically used as a loop diuretic in patients with edema and hypertension^54^ due to its inhibition of NKCC2 in renal tissue. However, bumetanide also inhibits NKCC1 which is expressed more systemically. This includes neurons where NKCC1 affects cell polarization and levels of GABA that are important for proper CNS function^91–93^. To that end, bumetanide has been used more recently as an anti-seizure medication in mice^92^ and off-label in humans^93^. In addition, oral solutions of bumetanide were recently used in clinical trials for autism in children^94^. However, bumetanide still impacts renal function when used for neurological symptoms, causing diuretic side effects^95^. These side effects are likely more challenging for those who are already medically fragile such as DPAGT1-CDG patients. However, the structural basis for bumetanide inhibition was recently described^54^, and this could pave the way for future bumetanide-like drugs. It is possible that such drugs could be used as anti-seizure medications without the diuretic side effects. Given that bumetanide improved the eye development of the *DPAGT1* model, it also suggests a role for bumetanide-like drugs beyond their use as anti-seizure medications.

The COX inhibitors tolmetin and antipyrine improved the *DPAGT1* model. While now discontinued in the US (via PubChem^96^), tolmetin was used for decades as an alternative to aspirin and has similarly few side effects^97^. Antipyrine, also known as phenazone, is an antipyretic currently used to treat ear infections^98^. In the past, antipyrine was used orally for decades and was one of the earliest prescribed analgesics^99^. Interestingly, while the antipyrine metabolite, 4-hydroxyantipyrine, was a hit in the screen, it did not validate in our hands. This could indicate some volatility of this metabolite or that a different dose is required. Inhibiting the cyclooxygenases COX-1 and COX-2 is a common drug mechanism for alleviating inflammation and pain^57,58^. However, there is also evidence for the use of COX-1 and COX-2 inhibitors in treating cognitive deficits and convulsions^59,100,101^, both of which are relevant to DPAGT1-CDG. While only two NSAIDs improved our model, this category may still be promising for DPAGT1-CDG given their high usage and ease of patient access^62^.

In this study, we screened 1,520 small molecules and identified multiple drugs that improved a *Drosophila* model of DPAGT1-CDG. We verify these findings using pharmacologic and genetic manipulation which strongly align with the known mechanisms of these therapeutic hits. We establish new biological connections between DPAGT1 and dopamine signaling, NKCC1, and prostaglandin synthesis. These findings may help create new treatment options for DPAGT1-CDG, and our validated drug hits represent potential therapeutics for patients.

## Materials and Methods

### Fly stocks and maintenance

All flies were maintained at room temperature. All experiments were performed in a 20°C incubator unless otherwise noted. Flies were fed standard Glucose medium (D2) from Archon Scientific (Durham, North Carolina). We used fly stocks from the Bloomington Drosophila Stock Center (NIH P40OD018537) and the Vienna Drosophila Resource Center^102^ (listed in S3 Table). The *DPAGT1* model (*eya* composite-GAL4, UAS-*Alg7* RNAi [III]) was described previously^21^. Its genetic background control, *eya* composite-GAL4 (III), was a gift from Justin Kumar (Indiana University Bloomington).

The *DPAGT1* model containing the *Dop1R1* null mutant (BDSC 92640) recombination (*DPAGT1* model [*Dop1R1^+/-^*]) was generated as follows. We crossed the *DPAGT1* model^21^ to the *Dop1R1* null mutant (BDSC 92640) to create (*eya composite*-GAL4, w+, P[sc+, y+, v+, *Alg7* RNAi])/(*Dop1R1* null) (III) animals. We then crossed this line to a balancer line containing *D*/(TM3, *ser*) (III) and examined progeny for crossover events. We collected progeny with smaller eye size than the *DPAGT1* model and *ser*. We then replaced the (TM3, *ser*) balancer with (TM3, *Sb*), self-crossed the line to ensure stability of the phenotype, and refer to this line as *DPAGT1* model [*Dop1R1^+/-^*] flies (*eya composite*-GAL4, w+, P[sc+, y+, v+, *Alg7* RNAi], *Dop1R1* null)/(TM3*, Sb*) (III).

### *in vivo* small molecule screen

For the primary screen, we used the Prestwick Chemical Library (“PCL”, Illkirch, France). The PCL contains 19 plates of 80 compounds each (1,520 total). All compounds came dissolved in DMSO at a concentration of 10 mM. Compounds were diluted further to 1 mM in phosphate-buffered saline. To make food containing these compounds, we used 500cc bags of standard Glucose medium (D2) from Archon Scientific (Durham, North Carolina). We dispensed this media into a beaker, microwaved until it liquified, then allowed it to cool on a heated stir plate under agitation. Once reaching 60°C, 1 mL of this media was dispensed into vials containing aliquots of the dissolved drugs or DMSO to reach a final concentration of 5 μM and 0.05% DMSO. We used up to eight DMSO control vials for each set of 80 drug vials.

Once the food was cooled, we placed 3-5 premated *DPAGT1* model intercrossed females and 2-3 males into each vial. Flies were allowed to lay eggs for 1-4 days until visual inspection indicated that approximately 30 eggs were laid. Flies were then removed. The *DPAGT1* model is balanced by TM3, *Sb*^21^, and homozygotes are infrequent and semi-lethal. As such, we only collected heterozygote flies by selecting for the *Sb* phenotype. We collected up to five (average = 4.4, males and females), 2-7 day old progeny flies from each vial and froze them at -80°C for later imaging. Drug names were masked, so both preparation of vials and collection of flies was done blinded to each drug.

Progeny eyes were imaged at 3x magnification (Leica EC3 camera). We determined eye area as previously described^103^. We masked image file names to blind observers to each treatment used. Within each set of 80 vials, we compared drug-treated animal eye sizes to the average of up to 8 DMSO controls and determined Z-scores (S1 Table). In one plate, drug-treated animals were instead compared to the plate average due to a technical error in the control (plate 03, S1 Table), but these values were in line with the rest of the plates. To determine if a drug was a hit, we used a Z-score threshold of 1.5 (derived from other *Drosophila* screens^104–106^).

We excluded all drug vials with a positive Z-score that contained only one fly (13 total vials, 0.9%). 15 compounds were excluded for technical reasons (e.g. incorrect food pours, 1%). An additional 16 vials had eggs laid without any eclosed flies (1.1%). If one or less males were observed, but females were present, females were measured instead; this occurred in 47 vials (3.1%), and none of these reached our Z-score threshold.

### Drug and RNAi validation

For making drug validation food, we used the same method as the primary screen, except that we used 10 mL of media, and we used drug concentrations of 1 μM, 5 μM, and 25 μM. Because there may be slight differences in drug concentrations from vendors or during food preparation, we considered drugs to validate if any of the three concentrations improved eye size (after multiple comparison correction). Most compounds were dissolved in DMSO. However, we employed other solvents if needed to reach the maximum concentration of 25 μM (S2 Table). We validated using at least two compounds from each drug category (S2 Table) from secondary vendors to limit quality control errors. These vendors were either Cayman Chemical (Ann Arbor, MI), MedChemExpress (Monmouth Junction, NJ), or Sigma-Aldrich (St. Louis, MO). Specific compounds and their vehicles are listed in S2 Table.

We crossed each fly knockout, overexpression, or RNAi line to either the *DPAGT1* model or its control, *eya* composite-GAL4 (S3 Table). All RNAi in this study is driven by the *eya* composite-GAL4 line. RNAi or overexpression crosses used either attP40 (BDSC 36304), attP2 (BDSC 36303), or attP (VDRC 60100) as control comparisons, and null crosses used *w*^1118^ (VDRC 60000). Because the *DPAGT1* model uses RNAi endogenously, RNAi crosses result in double knockdown flies. When fly stocks were available, we used two different RNAi lines to reduce potential reagent-specific effects. Complete information on lines used can be found in S3 Table. The average N across every drug validation experiment was 14.5 flies (based on 3,186 fly measurements).

Both drug and RNAi validation used the same methodology as the primary screen for collecting animals and measuring eye sizes.

### Statistics

For group comparisons, eye sizes were analyzed by One-way ANOVA with Welch’s and Dunnett’s corrections. For individual comparisons, eye sizes were analyzed by Student’s t test with Welch’s correction. We used GraphPad Prism v10 or Microsoft Excel for these analyses.

## Acknowledgements

We thank Emily Coelho for technical assistance with fly management.

